# Distinct effects of intracellular vs. extracellular acidic pH on the cardiac metabolome during ischemia and reperfusion

**DOI:** 10.1101/2022.08.17.504284

**Authors:** Alexander S. Milliken, Jessica H. Ciesla, Sergiy M. Nadtochiy, Paul S. Brookes

## Abstract

Tissue ischemia results in intracellular pH (pH_IN_) acidification, and while accumulation of metabolites such as lactate is a known driver of acidic pH_IN_, less is known about how acidic pH_IN_ regulates metabolism. Furthermore, acidic extracellular (pH_EX_) during early reperfusion confers cardioprotection, but how this impacts metabolism is unclear. Herein we employed LCMS based targeted metabolomics to analyze perfused mouse hearts exposed to: (i) control perfusion, (ii) hypoxia, (iii) ischemia, (iv) enforced acidic pH_IN_, (v) control reperfusion, and (vi) acidic pH_EX_ (6.8) reperfusion. Surprisingly little overlap was seen between metabolic changes induced by hypoxia, ischemia, and acidic pH_IN_. Acidic pH_IN_ elevated metabolites in the top half of glycolysis, and enhanced glutathione redox state. Acidic pH_EX_ reperfusion induced substantial metabolic changes in addition to those seen in control reperfusion. This included elevated metabolites in the top half of glycolysis, prevention of purine nucleotide loss, and an enhancement in glutathione redox state. These data led to parallel hypotheses regarding potential roles for methylglyoxal inhibiting the mitochondrial permeability transition pore, and for acidic inhibition of ecto-5’-nucleotidase, as potential mediators of cardioprotection by acidic pH_EX_ reperfusion. However, neither hypothesis was supported by subsequent experiments. In contrast, analysis of cardiac effluents revealed complex effects of pH_EX_ on metabolite transport, suggesting that mildly acidic pH_EX_ may protect in part by enhancing succinate release during reperfusion. Overall, each intervention had distinct and overlapping metabolic effects, suggesting acidic pH is an independent metabolic regulator regardless which side of the cell membrane it is imposed.

**HIGHLIGHTS:**

- Hypoxia, ischemia and acidic pH_IN_ each induce unique cardiac metabolic profiles.
- Acidic pH_EX_ at reperfusion prevents purine loss and enhances succinate release.

## 1. INTRODUCTION

pH is arguably one of the most fundamental environmental conditions that govern cell fate (along with factors such as temperature, oxygen and substrate availability). The primary metabolic sources of H^+^ in cells include lactic acid from glycolysis and CO_2_ from the TCA cycle [1]. Along with enzymes such as carbonic anhydrase [2], cells express a variety of transporters and exchangers at the plasma membrane to regulate intracellular pH (pH_IN_). This includes Na^+^/H^+^ exchangers (NHEs) [3], Na^+^/HCO_3_^-^ cotransporters (NBCs) [4, 5], and monocarboxylate transporters (MCTs) [6, 7] (**Figure 1A**).

**Figure 1.**
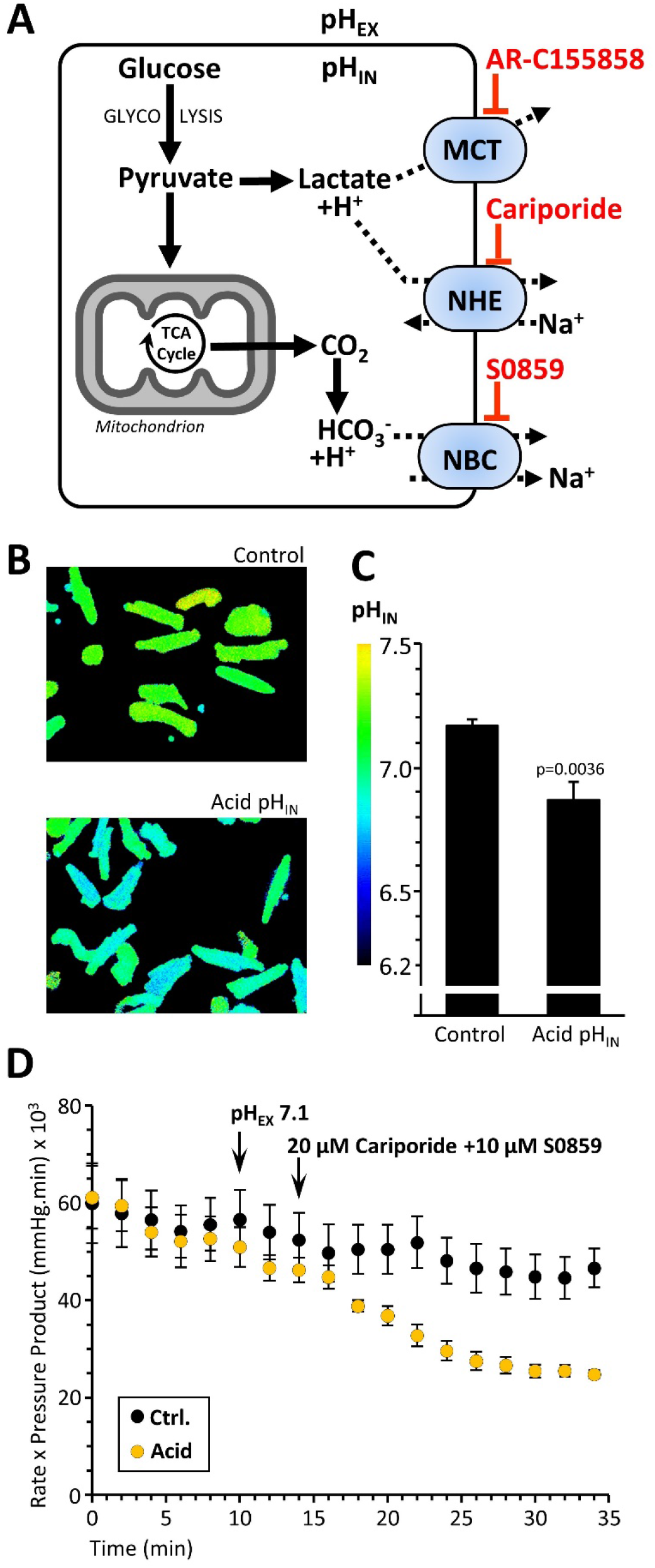
Cardiac pH_IN_ Manipulation. **(A):** Endogenous regulation of cardiomyocyte intracellular pH (pH_IN_). Glycolysis generates pyruvate, which can either be shuttled into mitochondria for TCA cycle activity (generating CO_2_) or remain in the cytosol to generate lactate and a proton (H^+^). Cardiomyocytes regulate intracellular pH via monocarboxylate transporters (MCTs), Na^+^/H^+^ exchangers (NHEs), and Na^+^/HCO^3-^ cotransporters (NBCs). Pharmacologic inhibitors for these exchangers and transporters used herein are shown in red: AR-C155858 (MCT inhibitor), cariporide (NHE1 inhibitor), and S0859 (NBC inhibitor). **(B):** Representative images of isolated cardiomyocytes subject to control and enforced acid pH_IN_ (cariporide + S0859 + pH_EX_ 7.1). **(C):** Quantitation of pH_IN_. Data are means + SEM, N = 4-5. p-value (Tukey’s HSD) is reported above error bar. Other conditions tested are shown in **Figure S2**. **(D):** Graph showing heart rate x left ventricular developed pressure (i.e., rate pressure product, RPP) of control and enforced acid pH_IN_ Langendorff perfused mouse hearts. Data are means ± SEM (N=8). Data for individual components (heart rate and developed pressure) are in **Figure S3**.

Despite classical knowledge that nearly all metabolic enzymes are pH sensitive, only recently has acidic pH been recognized as a metabolic remodeling signal. Specific examples include an acid-induced neomorphic enzyme activity in lactate and malate dehydrogenases, to generate the metabolic side-product *L*-2-hydroxyglutarate [8, 9], and an acid-induced reversal of isocitrate dehydrogenase to reductively carboxylate α-ketoglutarate to citrate [8, 10]. Both these metabolic events are thought to be characteristic of and advantageous to cancer cells [11]. The expulsion of H^+^ from cells also serves important signaling roles such as maintaining the tumor microenvironment [12], or in the case of stem cells maintaining pluripotency/stemness [13]. Ligand binding to cell surface receptors [14] and metabolite dynamics (i.e. import and export) [15] are also known to be pH sensitive.

The role of pH in determining outcomes in pathogenic states such as cardiac ischemia (i.e., cessation of coronary blood flow) is not fully understood. During cardiac ischemia, mitochondrial oxidative phosphorylation (Ox-Phos) is inhibited, and anaerobic glycolysis serves as a primary source of ATP [16], resulting in pH_IN_ acidification via lactate production. Similar effects are also seen during hypoxia (i.e., limited oxygen availability but with retention of substrate delivery and blood flow). Although ischemia and hypoxia have several overlapping features, the degree to which pH_IN_ acidification which occurs in both these conditions can act as an independent factor that remodels metabolism, is unclear. Herein we applied targeted metabolomics, to compare the effects of hypoxia, ischemia or enforced pH_IN_ acidosis on cardiac metabolism. We termed this study “CHIA” (<u>c</u>ontrol, <u>h</u>ypoxia, <u>i</u>schemia, <u>a</u>cid pH_IN_).

Following a cardiac ischemic event, paradoxically much of the subsequent pathology is triggered by tissue reperfusion (re-establishment of coronary blood flow), leading to ischemia-reperfusion (IR) injury. Upon reperfusion, mitochondrial Ca^2+^ overload and a burst of reactive oxygen species (ROS) generation combine to trigger opening of the mitochondrial permeability transition (PT) pore, leading to necrosis [17–21]. The major source of ROS at reperfusion is the rapid mitochondrial oxidation of succinate, which accumulates during ischemia [22–24]. However, around ⅔ of accumulated succinate washes out from the heart upon reperfusion, in a pH-sensitive manner. An acidic extracellular pH (pH_EX_ 6.0) can promote retention of succinate inside cells [15], suggesting that the local pH environment in the heart at reperfusion could influence succinate-driven ROS and IR injury [25].

Further highlighting the importance of pH during early reperfusion, extending ischemic acidosis into reperfusion via several independent methods has been demonstrated to confer cardioprotection. This includes: (i) reperfusion of the heart for 2 minutes with mildly acidic extracellular media (pH_EX_ 6.8) [26, 27], (ii) pharmacologic inhibition of carbonic anhydrase [28, 29], (iii) acute stimulation of glycolytic lactate production via bolus delivery of the NAD^+^ precursor NMN [30], and (iv) ischemic post-conditioning (aka, staccato reperfusion) [31]. In addition, although inhibition of NHEs confers cardioprotection [32], this effect has been mostly attributed to preventing myocardial Na^+^ overload, with effects on pH_IN_ largely overlooked.

Although acidic pH_IN_ is a known inhibitor of PT pore opening [33, 34], other downstream mediators of protection by acidic pH_EX_ reperfusion are not fully understood. Furthermore, an apparent paradox exists, in which acidic pH_EX_ at reperfusion is cardioprotective, but it should also drive succinate retention in cells (vide supra), which would lead to more ROS generation and damage. Thus, herein we applied metabolomics analysis to examine the impact of acidic pH_EX_ during early reperfusion on cardiac metabolism. We termed this study “A-pH_EX_”. We also explored several downstream metabolic mechanisms as potential mediators of acidic pH_EX_ induced cardioprotection. Overall, in addition to providing reference data sets for the effects of pH on cardiac metabolism, our results indicate that acidic pH_IN_ and pH_EX_ are powerful metabolic regulators with both overlapping and independent effects.

## 2. MATERIALS AND METHODS

### 2.1 Animals and Reagents

Animal experiments complied with the NIH *Guide for Care and Use of Laboratory Animals* (8^th^ edition, 2011) and were approved by the University of Rochester Committee on Animal Resources. Male and female C57BL/6J mice (8-16 weeks old) were housed on a 12 hr. light/dark cycle with food and water *ad libitum*. Terminal anesthesia was achieved via intra-peritoneal 2,2,2-tribromoethanol (Avertin) at 250 mg/kg. BCECF-AM was from Molecular Probes (Eugene, OR, USA) and collagenase was from Roche (Indianapolis, IN, USA). All other reagents were from Sigma (St. Louis, MO, USA) or MedChemExpress (Monmouth Jct., NJ, USA).

### 2.2 Adult Mouse Cardiomyocyte Isolation

Following anesthesia, the aorta was rapidly cannulated, then the heart excised and perfused for 3 min. with Isolation Buffer (IB), comprising (in mM): NaCl (120), KCl (15), Na_2_HPO_4_ (0.6), KH_2_PO_4_ (0.6), MgSO_4_ (1.2), HEPES (10), NaHCO_3_ (4.6), taurine (30), glucose (5.5), butanedione monoxime (10), pH 7.4 at 37 °C. The heart was then perfused for 10 min. with Digestion Buffer (DB), comprising IB plus 12.5 μM CaCl_2_, 0.025 % (wt/vol) trypsin, 6.525 U collagenase A, and 15.375 U collagenase D. Ventricular tissue was teased apart, resuspended in Stop Buffer (SB), consisting of IB plus 12.5 μM CaCl_2_ and 10 % (vol/vol) FBS, and filtered through 200 μm mesh. Cells were settled by gravity for 10 min., then sequentially suspended/settled in SB with [Ca^2+^] increased stepwise to 1.8 mM. Finally, cells were suspended in DMEM with (in mM) L-glutamine (4), Na-pyruvate (0.1), glucose (5), L-carnitine (0.5), palmitate conjugated to BSA (0.1), pH 7.4 at 37 °C. This protocol [35] yielded ^~^8 x 10^5^ cells per heart, with ^~^80% rod-shaped cells excluding Trypan blue dye.

### 2.3 Measurement of cardiomyocyte pH_IN_

Cardiomyocytes were seeded on laminin-coated 35 mm coverslips (20 μg/ml) for 1 hr., incubated with the pH-sensitive dye 2’-7’-bis-(2-carboxyethyl)-5-(and-6)-carboxyfluorescein, acetoxymethyl ester (BCECF-AM) at 1.26 mM for 30 min., then subject to various combinations of the following treatments for 1 hr.: (i) Control pH 7.4, (ii) pH_EX_ 7.1, (iii) 100 nM of the protonophore carbonyl cyanide-p-trifluoromethoxyphenylhydrazone (FCCP), (iv) 30 μM of the NBC inhibitor S0859, (v) 20 μM of the NHE1 inhibitor cariporide, (vi) 1 mM of the NAD^+^ precursor nicotinamide mononucleotide (NMN), (vii) 1 μM of the MCT-1 inhibitor AR-C155858. Ratiometric fluorescence was measured at λ_EX_ /λ_EM_ 440/535 and 440/490 nm using an Eclipse TE2000-S microscope (Nikon, Avon MA) and data were analyzed using a TILL Photonics System as previously described [8]. pH_IN_ was calculated from a determined Boltzmann factor calibration curve using the ionophore nigericin based on fluorescence ratios. For each coverslip, four images were taken, capturing 5-16 individual cardiomyocytes (technical replicates) to determine pH_IN_ values. Groups of 3-4 coverslips on a given day were considered as biological replicates (N).

### 2.4 Langendorff-perfused mouse hearts

Following anesthesia, the aorta was rapidly cannulated, the heart excised and perfused at 4 ml/min. with Krebs-Henseleit buffer (KHB) comprising (in mM): NaCl (118), KCl (4.7), MgSO_4_ (1.2), NaHCO_3_ (25), KH_2_PO_4_ (1.2), CaCl_2_ (2.5), glucose (5), pyruvate (0.2), lactate (1.2), palmitate conjugated 6:1 with fat-free BSA (0.1). KHB was gassed with 95 % O_2_ and 5 % CO_2_ at 37 °C to maintain pH 7.4. A water-filled balloon connected to a pressure transducer was inserted into the left ventricle and expanded to provide a diastolic pressure of 6-8 mmHg. Cardiac function was recorded digitally at 1 kHz (Dataq, Akron OH).

To determine the effect of hypoxia, ischemia, or acidic pH_IN_ on cardiac metabolism, hearts were equilibrated for 15 min., and then further subjected to 20 min. of either: (i) control perfusion, (ii) hypoxia (buffer gassed with 95% N_2_, 5% CO_2_), (iii) global no-flow ischemia, or (iv) acidic pH_IN_ (**Figure 2A**). Optimal acidic pH_IN_ conditions (20 μM cariporide + 10 μM S0859, at pH_EX_ 7.1) were determined from cardiomyocyte experiments, see **Figure S1**. This study is referred to as “CHIA” (<u>c</u>ontrol, <u>h</u>ypoxia, <u>i</u>schemia, <u>a</u>cid pH_IN_).

**Figure 2.**
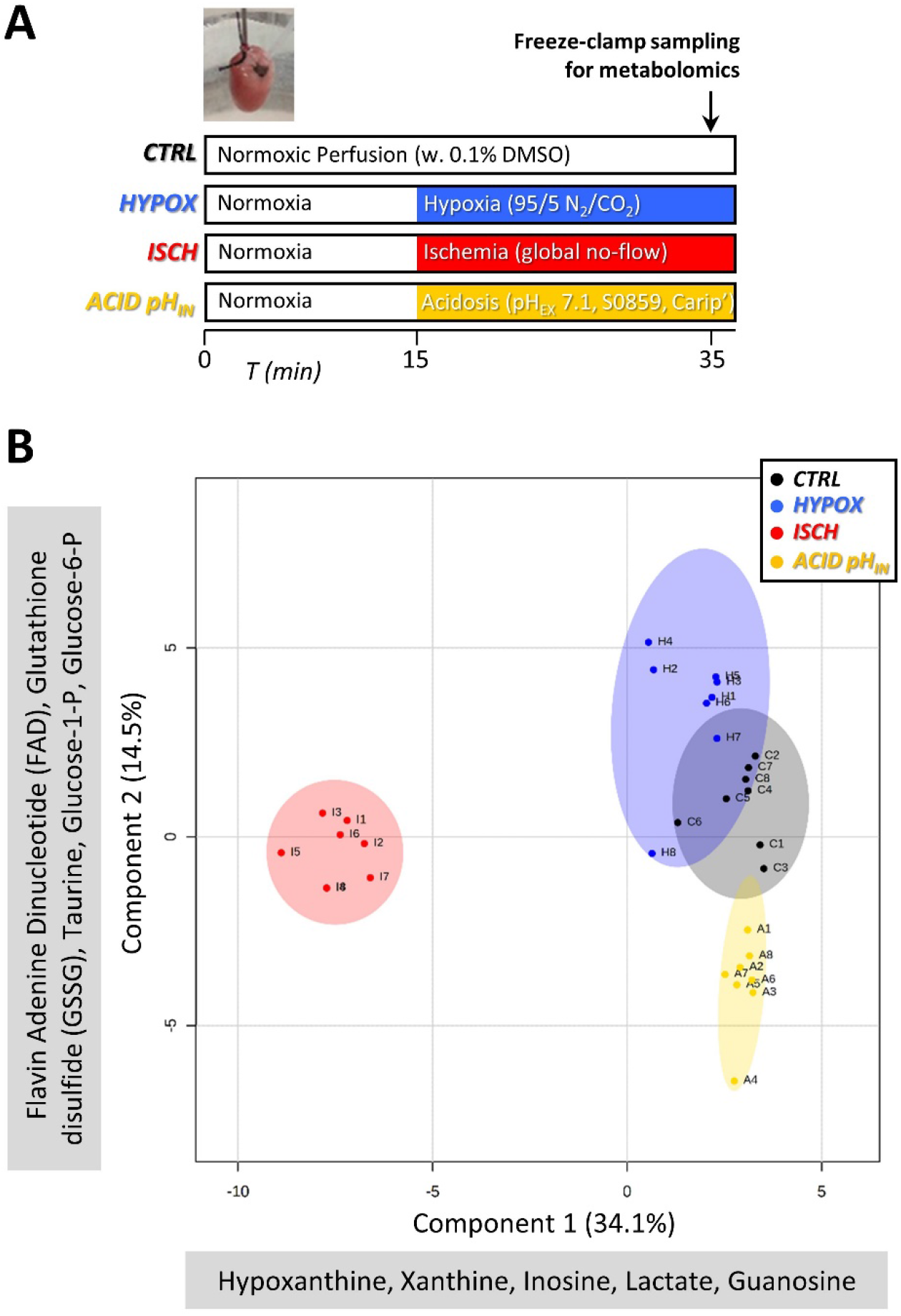
Metabolomics CHIA study (<u>C</u>ontrol, <u>H</u>ypoxia, <u>I</u>schemia, <u>A</u>cidosis). **(A):** Schematic showing design of the 4 experimental perfusion conditions in the CHIA study. Hearts were snap frozen for steady-state metabolomics analysis by LC-MS indicated by the arrow. Colors and terms used to denote each condition are used throughout. **(B):** Sparse partial least-squares discriminant analysis (sPLSDA) plot, prepared using MetaboAnalyst 5.0, incorporating 59 metabolites from 32 samples. Key at upper right denotes symbols for each group. The top 5 metabolite weightings contributing to each principal component (Component 1: x-axis, Component 2: y-axis) are shown alongside each axis.

To examine the impact of acidic pH_EX_ reperfusion on cardiac metabolism, following a 15 min. equilibration hearts were either (i) sampled immediately, or (ii) after being further subjected to 25 min. global no-flow ischemia, or (iii) 25 min. global no-flow ischemia plus 2 min. control reperfusion (pH_EX_ 7.4), or (iv) 25 min. global no-flow ischemia plus 2 min. acidic reperfusion (pH_EX_ 6.8) (**Figure 4A**). This study is hereafter referred to as “A-pH_EX_”.

**Figure 3.**
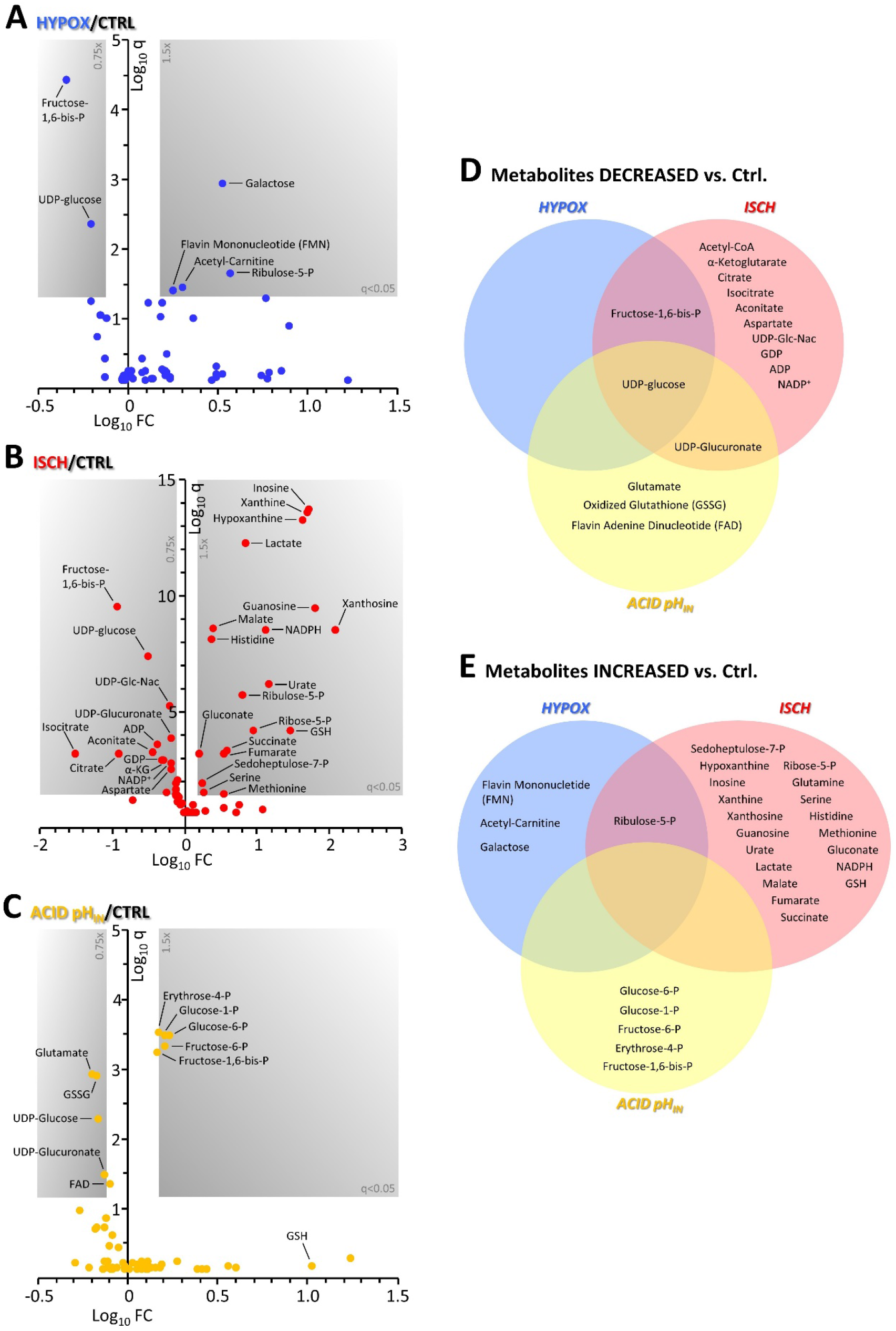
Metabolomics reveals differential effects of hypoxia, ischemia, acidosis. Volcano plots showing relative metabolite abundance in **(A)** hypoxia, **(B)** ischemia, and **(C)** acid pH_IN_, vs. control perfusion. Log_10_ fold changes are on x-axes and Log_10_ FDR-corrected p-values (q-values) on y-axes. Metabolites in gray shaded areas are significantly (q-value < 0.05) up or down (>1.5 fold). Data are means, N=8 per condition. **(D/E):** Venn diagrams showing commonalities and differences between metabolites either decreased **(D)** or increased **(E)** by each of the 3 perturbations.

**Figure 4.**
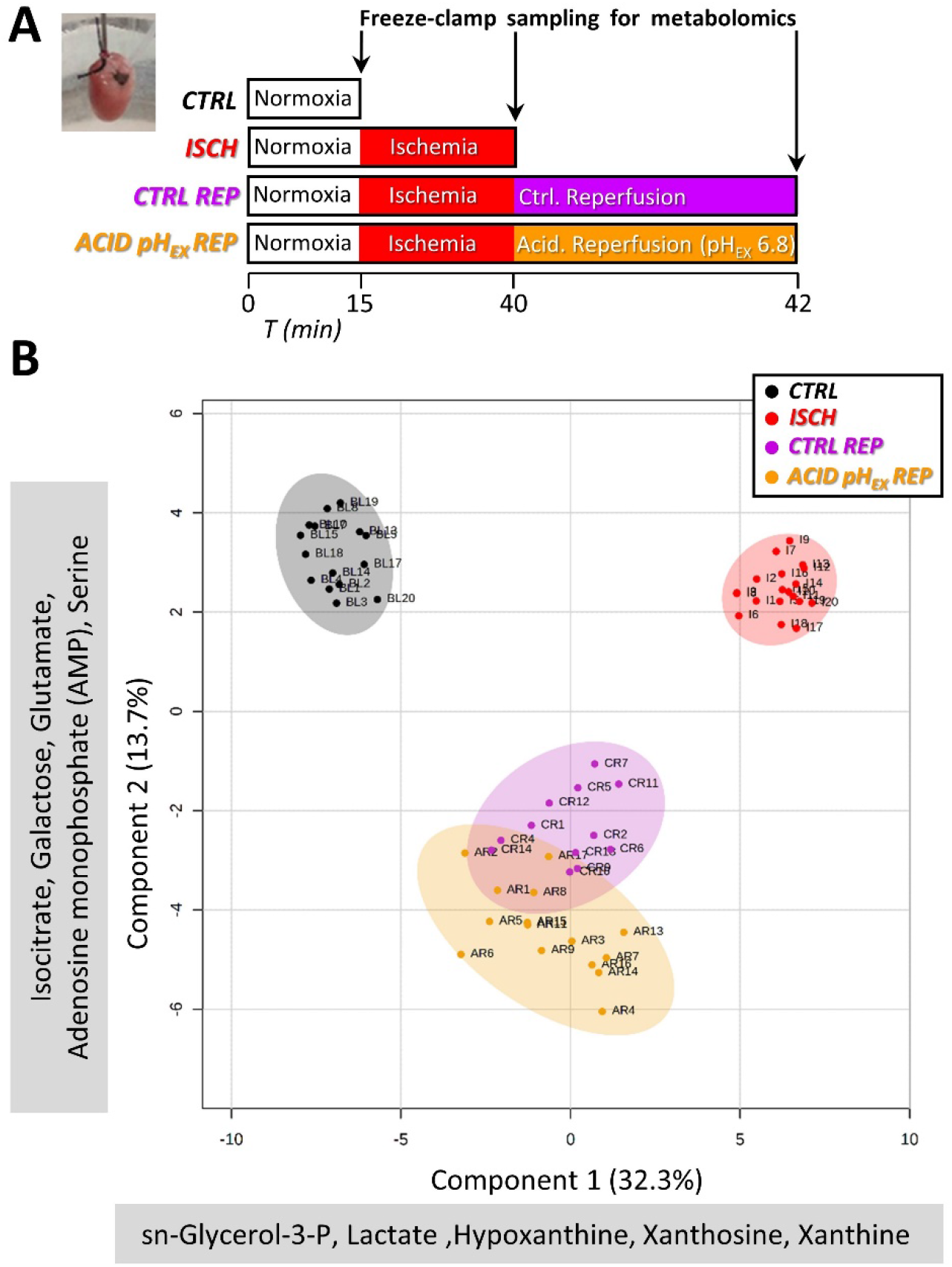
Metabolomics, A-pH_EX_ study (acidic pH_EX_ reperfusion). **(A):** Schematic showing design of the 4 experimental perfusion conditions in the A-H_EX_ study. Hearts were snap frozen for steady-state metabolomics analysis by LC-MS as indicated by the arrows. Colors and terms used to denote each condition are used throughout. **(B):** Sparse partial least-squares discriminant analysis (sPLSDA) plot, prepared using MetaboAnalyst 5.0, incorporating 79 metabolites from 61 samples. Key at upper right denotes symbols for each group. The top 5 metabolite weightings contributing to each principal component (Component 1: x-axis, Component 2: y-axis) are shown alongside each axis.

Acidic pH_EX_ was obtained with a modified KHB containing 3.8 mM NaHCO_3_, yielding pH_EX_ 6.8 upon gassing with 95 % O_2_, 5 % CO_2_. During reperfusion, cardiac effluents were collected at 1 min. intervals for 2 min. Samples were immediately treated with 10 % perchloric acid, followed by addition of 100 nmols butyrate as internal standard, and stored at −80 °C for analysis.

To examine the effect of inhibiting ecto-5’-nucleotidase (5NT), hearts were subject to 25 min. global no-flow ischemia followed by 60 min. reperfusion, with optional infusion of 10 μM adenosine 5’-(α,β-methylene) diphosphate (meth-ADP), for 10 min. pre-ischemia and 2 min. post-ischemia into reperfusion.

For metabolomics, sampling comprised clamping the heart with pre-cooled Wollenberger tongs, plunging into liquid N_2_ and grinding to powder, followed by storage under N_2_ at-80 °C until analysis. Post-reperfusion hearts not used for metabolomics were transverse sliced (2 mm thick), stained with 1 % tetrazolium chloride, formalin fixed, and imaged for infarct measurement. Live (red) versus infarcted (white) tissue was quantified by planimetry using a custom MATLAB script.

### 2.5 Metabolite Analysis

Metabolites were serially extracted in 80 % methanol, evaporated under N_2_ then resuspended in 50 % methanol. Liquid Chromatography Tandem Mass Spectrometry (LC-MS/MS) analysis was performed as previously described [36], with reverse-phase LC on a Synergi Fusion-RP column (Phenomenex, Torrence CA) at 35 °C. Samples were analyzed by single reaction monitoring on a Thermo Quantum triple-quadrupole mass spectrometer. HPLC effluent was subject to electrospray ionization in negative ion mode using a HESI-II ion source. Metabolites were identified against a library of validated standards based on retention time, intact mass, collision energy, and fragment masses. Data were collected and analyzed using Thermo XCaliber 4.0 software. Post-hoc data analysis utilized Metaboanalyst 5.0 [37] for the generation of sPLSDA (sparse partial least-squares discriminant analysis) plots.

Each daily batch of 12 samples was bookmarked with a pooled sample to monitor instrument sensitivity. Following batch normalization, peaks were normalized to the sum of all peaks within each sample. Outliers were identified and discarded, as those values falling outside the 99.99 % confidence intervals for each group.

For the CHIA study, N for each group was 8. No samples were excluded, and across the 32 samples a total of 3.0 % of data were missing or discarded as outliers. A Jarque-Bera test for normality revealed 89 % of metabolites were normally distributed, permitting parametric statistical analysis. For the A-pH_EX_ study, the initial N for each group was: control 20, ischemia 20, normal reperfusion 14, acidic pH_EX_ reperfusion 17. Of these 71 original samples, 10 were excluded due to excessive noise (high numbers of outliers), and across the remaining 61 samples, a total of 3.3 % of data were missing or discarded as outliers. A Jarque-Bera test for normality revealed 84 % of metabolites were normally distributed.

Across the data sets for both CHIA and A-pH_EX_ studies, all missing values were imputed as medians of remaining values for each group [38]. Both studies contained control and ischemic groups, and although they were performed almost 3 years apart, and the latter study offered greater metabolome coverage (79 vs. 59 metabolites), comparison of the changes induced by ischemia in both studies revealed a significant correlation (r^2^ 0.83, **Figure S1**), suggesting that comparisons between them is permitted.

### 2.6 Effluent Analysis by High-Performance Liquid Chromatography

Effluents from control and acidic pH_EX_ reperfused hearts were analyzed by HPLC (Shimadzu Prominence 20A system) using two 300 x 7.8 mm Aminex HPX-87H columns (BioRad, Carlsbad CA, USA) in series, with 10 mM H_2_SO_4_ mobile phase (flow 0.7 ml/min) at 35 °C. 100 μl sample was injected. Carboxylic acids (succinate, fumarate, lactate) were quantified via photodiode array (A_210_) as previously described [39], with normalization to internal butyrate standard, and a standard curve for each metabolite.

### 2.7 Mitochondrial PT pore opening

Mouse liver mitochondria were isolated by differential centrifugation, and PT pore opening measured spectrophotometrically as osmotic swelling-induced decrease in light scatter/absorbance at 540 nm, both as previously described [36]. MGO or the pore inhibitor cyclosporin A (CsA) were added at concentrations indicated in figure legends, prior to triggering of PT pore opening by 100 μM Ca^2+^.

### 2.8 Western blotting

Heart or cardiomyocyte samples were separated by SDS-PAGE and immunoblotted to nitrocellulose, then probed with a mouse monoclonal anti-MGO adduct antibody (AbCam ab243074). Blot detection used secondary antibodies, ECL reagents, and a KwikQuant imaging system, all from Kindle Bioscience.

### 2.9 Statistical Analysis

Student’s t-test (paired and unpaired) and ANOVA with post-hoc Tukey’s HSD test were applied where appropriate. Significance was defined as α=0.05. For metabolomics data from the CHIA study, the Storey modification of the Benjamini-Hochberg correction for false discovery rate (FDR) was applied [40], with q-values (FDR-corrected p-values) displayed in results. For the A-pH_EX_ study, application of FDR correction revealed all metabolites exhibiting a significant p-value (Tukey’s HSD test) remained significant after FDR correction, so p-values are displayed in results.

## 3. RESULTS

The complete original data set is available on the data sharing site FigShare (DOI: 10.6084/m9.figshare.16602281 – DOI reserved, unembargoed upon publication).

### 3.1 Cardiomyocyte pH_IN_

Before proceeding to heart perfusions, we first screened a range of pharmacologic and physiologic treatments to induce pH_IN_ acidification in isolated primary adult mouse cardiomyocytes. Although cardiomyocytes are only 20 % of cells in the heart by number, they occupy 80 % of cardiac volume, suggesting treatment regimes validated in isolated cardiomyocytes could be applied to achieve acidification of pH_IN_ in the intact heart. As shown in **Figures 1B/C and Figure S2**, the combination of cariporide (NHE1 inhibitor), S0859 (NBC inhibitor) and pH_EX_ 7.1 yielded a drop in pH_IN_ of ^~^0.3 units, to pH 6.87. Although a similar degree of acidification was observed with MCT-1 inhibition (AR-C155858), we chose not to pursue this strategy due to its known effects on key metabolic pathways (lactate/succinate dynamics).

### 3.2 Acidic pH_IN_ lowers but does not stop cardiac function

**Figure 1D** shows that hearts treated with the combination of cariporide/S0859/pH_EX_ 7.1, exhibited a gradual decline in contractile function (rate x pressure product), stabilizing at around 55% of that seen in controls. The decline was driven at first by a lowering of heart rate (chronotropy) followed by a lowering of left-ventricular developed pressure (inotropy) (**Figure S3**). A fall in coronary root pressure was also seen (data not shown), consistent with the well-known vasodilatory effects of acid.

### 3.3 Control, hypoxia, ischemia, acidosis (CHIA) effects on cardiac metabolism

A total of 59 metabolites were reliably measured across all 32 samples examined in the CHIA study (N=8 per condition). Application of sparse partial least squares discriminant analysis (sPLSDA, a dimensionality reduction tool, **Figure 2**) revealed that the 4 treatment conditions distributed into independent clusters along 2 axes. Differences between control and ischemia moved primarily along component 1 (x-axis), whereas differences induced by hypoxia or acidic pH_IN_ moved primarily along component 2 (y-axis). This suggests the fundamental character of metabolic changes bought about by ischemia is different than the other 2 conditions. This is borne out in volcano plots (**Figures 3A-C**), illustrating the magnitude and statistical significance of metabolic changes brought about by the 3 perturbations. Ischemia induced a far greater number of changes than did hypoxia or acidic pH_IN_. A potential confounding factor in these divergent effects is the absence of coronary flow in ischemia, versus persistent flow in the hypoxia and acid pH_IN_ conditions.

A convenient tool to visualize the different metabolic changes induced by the 3 perturbations is Venn diagrams, as shown in **Figures 3D/E**. Surprisingly, only 1 metabolite change was shared between all 3 perturbations – namely a decrease in the glycogen synthetic precursor UDP-glucose. This suggests that acidic pH_IN_ may be an important signal in stimulating glycogen synthesis during ischemia or hypoxia. It has been reported previously that acidosis directly stimulates hepatic glycogen synthesis [41]. In addition, although glycogen is an important source of glucose during ischemia [24], it is known glycogen turnover is significant in the ischemic heart [42], suggesting that acidic pH_IN_ during ischemia may be an important signal for maintaining glycogen synthesis under conditions of enhanced glycogen utilization.

Further examining metabolites that were decreased, a lowering of fructose-1,6,-bis-phosphate (F-1,6-BP) was seen in both hypoxia and ischemia, but notably this metabolite was elevated by acidic pH_IN_. This is somewhat counterintuitive, since acidic pH is known to inhibit PFK1, the enzyme that produces F-1,6-BP. Furthermore, several other metabolites in the “top-half” of glycolysis (i.e., glucose-6-P, glucose-1-P, fructose-6-P) were all significantly increased by acidic pH_IN_. This is consistent with an acid-induced inhibition at the level of GAPDH (the forward reaction of which generates a H^+^), and this mid-point of glycolysis is increasingly viewed as a key regulatory stage for the pathway [43, 44]. These differential effects of acid pH_IN_ alone, versus acidification that occurs in the context of ischemia or hypoxia, would appear to suggest that in the latter 2 conditions, other factors may over-ride the effect of acid on glycolysis. This could include limited substrate availability in the case of no-flow ischemia, or the need to maintain glycolytic flux to provide ATP in the absence of mitochondrial Ox-Phos. Overall, it appears that perturbations to glycolytic activity seen in ischemia are unlikely driven purely by the effects of acidic pH_IN_.

A number of other changes were observed in hearts subjected to acidic pH_IN_. In particular, a 32% drop in oxidized glutathione (GSSG), coupled with a 10-fold rise in reduced glutathione (GSH, although this latter change did not reach statistical significance) together suggest that acidic pH_IN_ may favor a more reduced cellular environment, enhancing resistance to oxidative stress. This is consistent with the general notion that acidic pH is cardioprotective [26–31]. Another notable feature of the acidic pH_IN_ metabolome was a depression in the levels of glutamate. Previously, we showed that anaplerosis into the TCA cycle from glutamate is a significant feature of ischemia [24], although the impact of acidic pH_IN_ on this was not considered. The current data suggest glutamate anaplerosis to α-ketoglutarate may be sensitive to by pH_IN_.

### 3.4 A-pH_EX_, the metabolic transition from ischemia to reperfusion

In addition to enforced pH_IN_, we examined the impact of acidic pH_EX_ for the first 2 minutes of reperfusion, because this regimen has been shown to induce cardioprotection. A total of 79 metabolites were reliably measured across 61 samples included in the A-pH_EX_ study (N=12-19 per condition). Application of sPLSDA (**Figure 4B**) revealed that the 4 treatment conditions distributed into independent clusters along 2 component axes. Similar to the CHIA study, differences between control and ischemia moved primarily along component 1 (x-axis), with several shared highly-weighted contributors between both studies (compare to **Figure 2B**). Upon reperfusion, both control and acidic pH_EX_ groups moved back along component 1 toward the control state, but also introduced novel features along a different axis, component 2 (y-axis). Notably, acidic pH_EX_ reperfusion diverged further along this axis than control reperfusion. Together these data suggest that control and acidic pH_EX_ reperfusion are largely similar in character, with the latter inducing greater changes.

**Figures 5A/B** show volcano plots for changes in metabolite levels with either control or acidic pH_EX_ reperfusion, compared to ischemia. **Figure 5C** shows key differences between metabolites in control or acidic pH_EX_ reperfused hearts. Visualization of these data as Venn diagrams (**Figures 5D/E**) indicates that the vast majority of metabolic changes induced by control reperfusion were also present in acidic pH_EX_ reperfusion (i.e., there were very few metabolic changes that were unique to control reperfusion and lost in acidic pH_EX_ reperfusion). In contrast, acidic pH_EX_ reperfusion induced a large number of additional metabolic changes that were unique to this condition and were not seen in control reperfusion.

**Figure 5.**
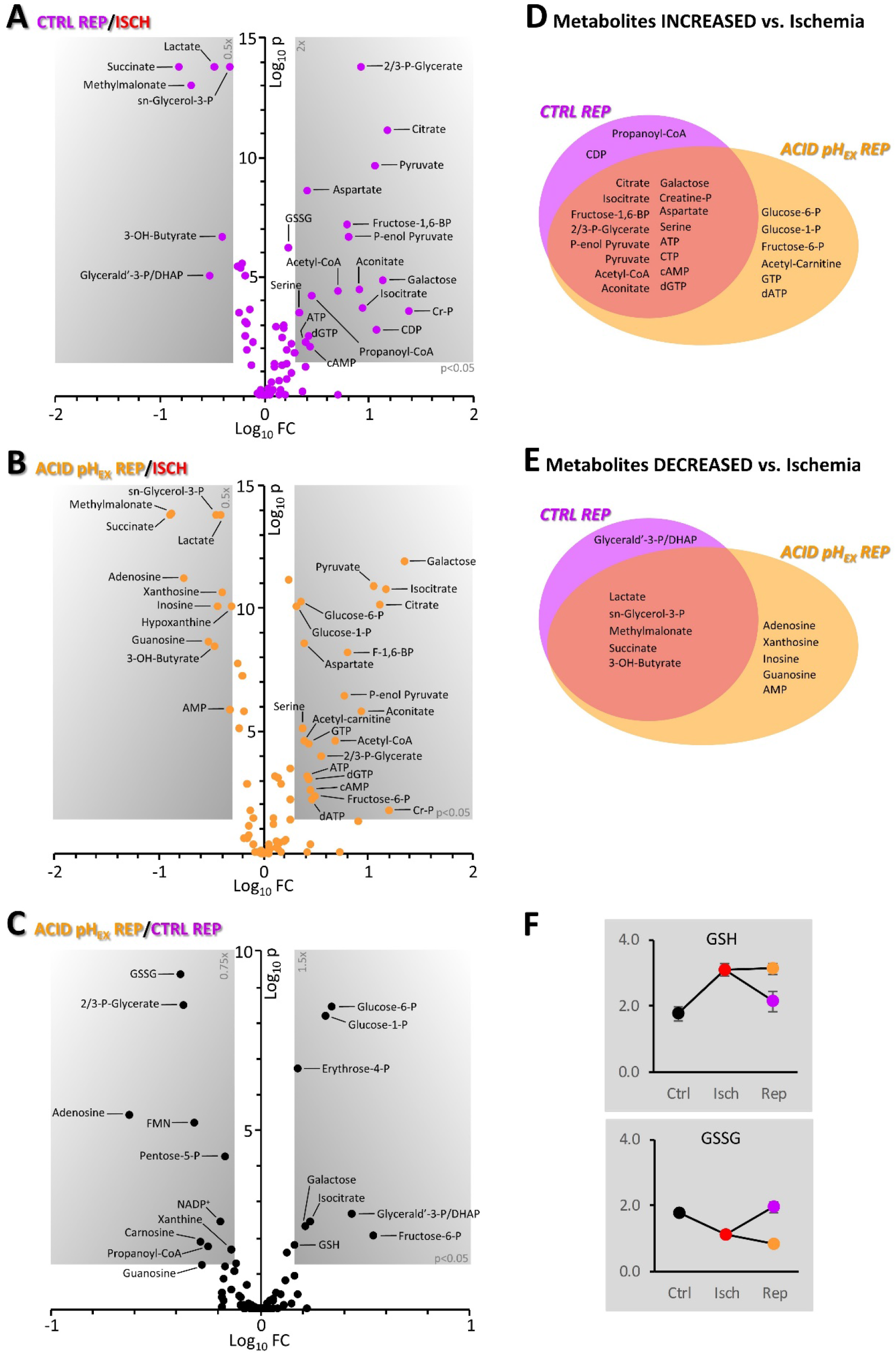
Metabolomics reveals differential effects of control vs. acid pH_EX_ reperfusion. **(A/B):** Volcano plots showing relative metabolite abundance transitioning from ischemia into either **(A)** control reperfusion (pH_EX_ 7.4) or **(B)** acid reperfusion (pH_EX_ 6.6). **(C):** Relative metabolite abundance comparing control and acid pH_EX_ reperfusion. For volcano plots, Log_10_ fold changes are on the x-axis and Log_10_ p-values are on the y-axis. Metabolites in gray boxes are significantly (p-value < 0.05) up or down (>2.0 fold). Data are means, N=12-19 per condition. All metabolites shown as significant remained so after Storey false discovery rate (FDR) correction. **(D/E):** Venn diagrams showing commonalities and differences between metabolites either decreased **(D)** or increased **(E)** as a result of either control or acid pH_EX_ reperfusion. **(F):** Relative changes in abundance of reduced and oxidized glutathione (GSH and GSSG respectively) from baseline into ischemia and then into either control or acid pH_EX_ reperfusion. Data are means ± SEM (N=12-19).

Metabolite changes upon reperfusion that were common to both conditions, were largely as expected. Most metabolites that decreased during ischemia were then increased upon reperfusion. This included several TCA cycle metabolites (citrate, isocitrate), as well as high energy phosphates (ATP, CTP, PCr) and metabolites in the lower half of glycolysis. Likewise, most metabolites that were elevated in ischemia were then decreased upon reperfusion in a manner that was not impacted by pH_EX_. This included succinate, lactate, and 3-OH-butyrate.

There were several metabolite differences between control and acidic pH_EX_ reperfusion hearts, including the redox state of glutathione. As **Figure 5F** shows, GSH was elevated and GSSG was lower in ischemia, and these changes were reversed upon control reperfusion. However, in acidic pH_EX_ reperfusion both GSH and GSSG were maintained at their ischemic levels into reperfusion. This suggests acidic pH_EX_ reperfusion may carry a significantly lower burden of oxidative stress than control. Although we have shown that cytosolic and mitochondrial pH can impact ROS generation by the respiratory chain [25], whether the cardioprotective benefits of acidic pH_EX_ reperfusion are conferred by a lowering of ROS generation, is currently unclear.

### 3.5 Cardioprotective roles for metabolites uniquely elevated by acidic pH_EX_ reperfusion

While most metabolites in the lower half of glycolysis returned to normal levels regardless of the reperfusion pH_EX_, this was not the case for the top half of the pathway. As shown in **Figure 6**, hexose phosphates were all significantly elevated with acidic pH_EX_ reperfusion. This is consistent with the elevation of these metabolites caused by acidic pH_IN_ in the CHIA study (**Figure 3**), and suggests that glycolysis responds rapidly to both pH_IN_ and pH_EX_.

**Figure 6.**
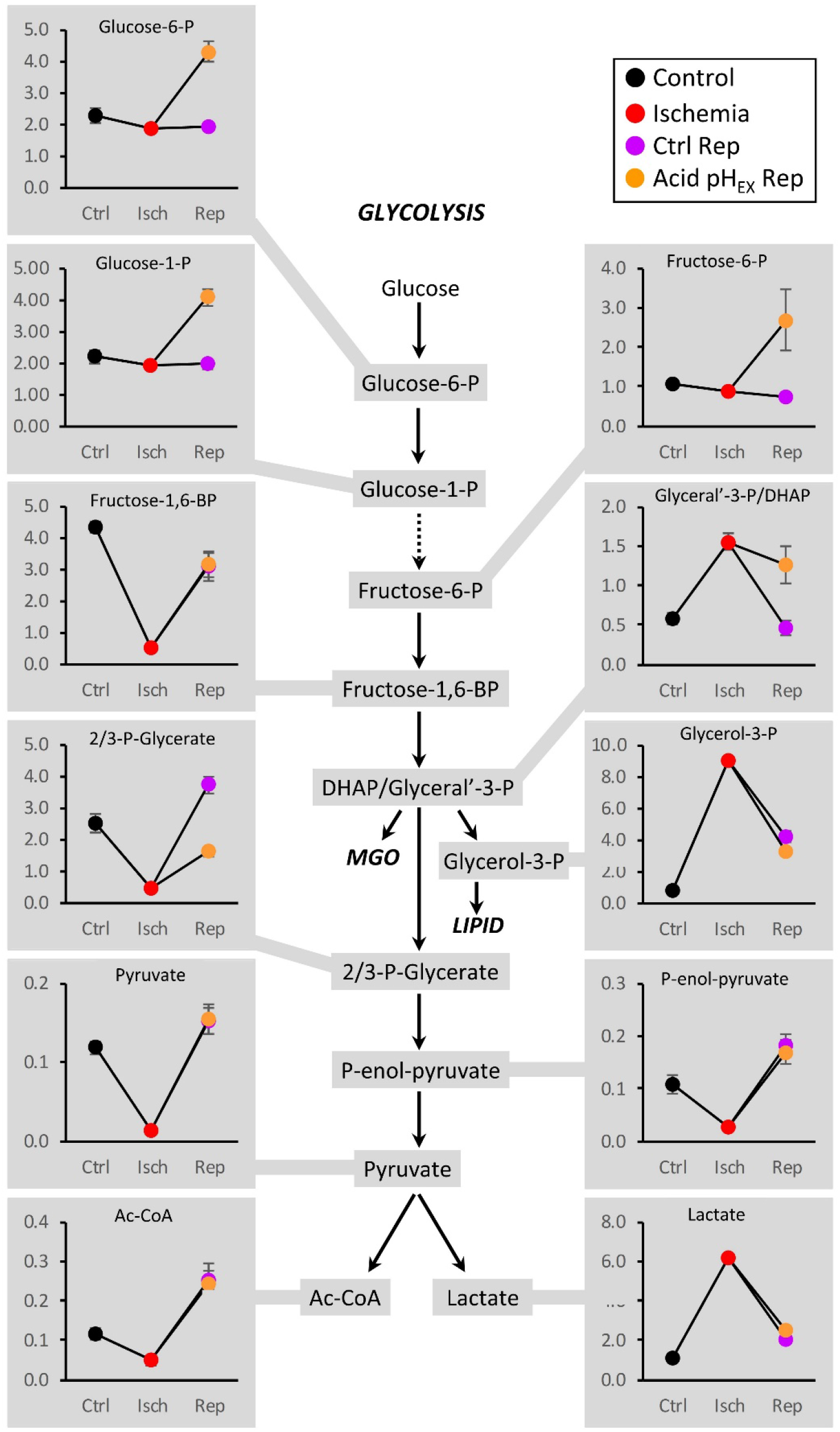
Metabolomics of glycolysis. The pathway is shown with each metabolite that was measured highlighted in gray and linked to its data. Key denotes the color scheme for each treatment group as used in previous figures. Data are means ± SEM (N=12-19 per condition) of relative metabolite abundance (arbitrary units).

A key observation was that levels of glyceraldehyde-3-phosphate / dihydroxyacetone-phosphate (DHAP) were retained in an elevated state upon acidic pH_EX_ reperfusion, versus declining with control reperfusion (**Figure 6**). An important byproduct generated from DHAP is the reactive metabolite methylglyoxal (MGO), which can post-translationally modify proteins (i.e., advanced glycation end products, AGEs) [45, 46]. It has also been shown that a related metabolite phenylglyoxal can inhibit the mitochondrial permeability transition pore [47, 48]. Given that acidic pH is a known inhibitor of the PT pore [33, 34], we hypothesized that MGO generation may be a molecular link between acidic pH and PT pore inhibition. As shown in **Figure S4**, MGO was able to inhibit PT pore opening in isolated mitochondria. However, western blotting for MGO protein adducts did not reveal any difference between control and acidic pH_EX_ reperfused hearts. Similar results were seen in primary cardiomyocytes exposed to acidic pH_EX_. Together these data suggest that MGO is unlikely responsible for the cardioprotection induced by acidic pH_EX_.

### 3.6 Cardioprotective roles for metabolites uniquely decreased by acidic pH_EX_ reperfusion

A number of differences were observed in the levels of purine nucleotides and their metabolites, between control and acid pH_EX_ reperfused hearts (**Figure 5E**). As shown in **Figure 7**, acid pH_EX_ reperfused hearts consistently had lower levels of nucleosides (guanosine, adenosine, inosine, xanthine, xanthosine, hypoxanthine) and elevated ratios of nucleotide triphosphates vs. diphosphates (GTP/GDP and ATP/ADP). This led us to hypothesize that the activity of ecto-5’-nucleotidase (5NT) may be inhibited by acidic pH_EX_, preventing purine nucleotide loss upon reperfusion. To determine whether this could play a role in the mechanism of cardioprotection elicited by acidic pH_EX_, we subjected hearts to IR in the presence of the 5NT inhibitor meth-ADP. However, as shown in **Figure S5**, this intervention failed to elicit cardioprotection. Thus, it appears that although acidic pH_EX_ reperfusion prevents purine nucleotide loss, this may not be responsible for the concomitant cardioprotection.

**Figure 7.**
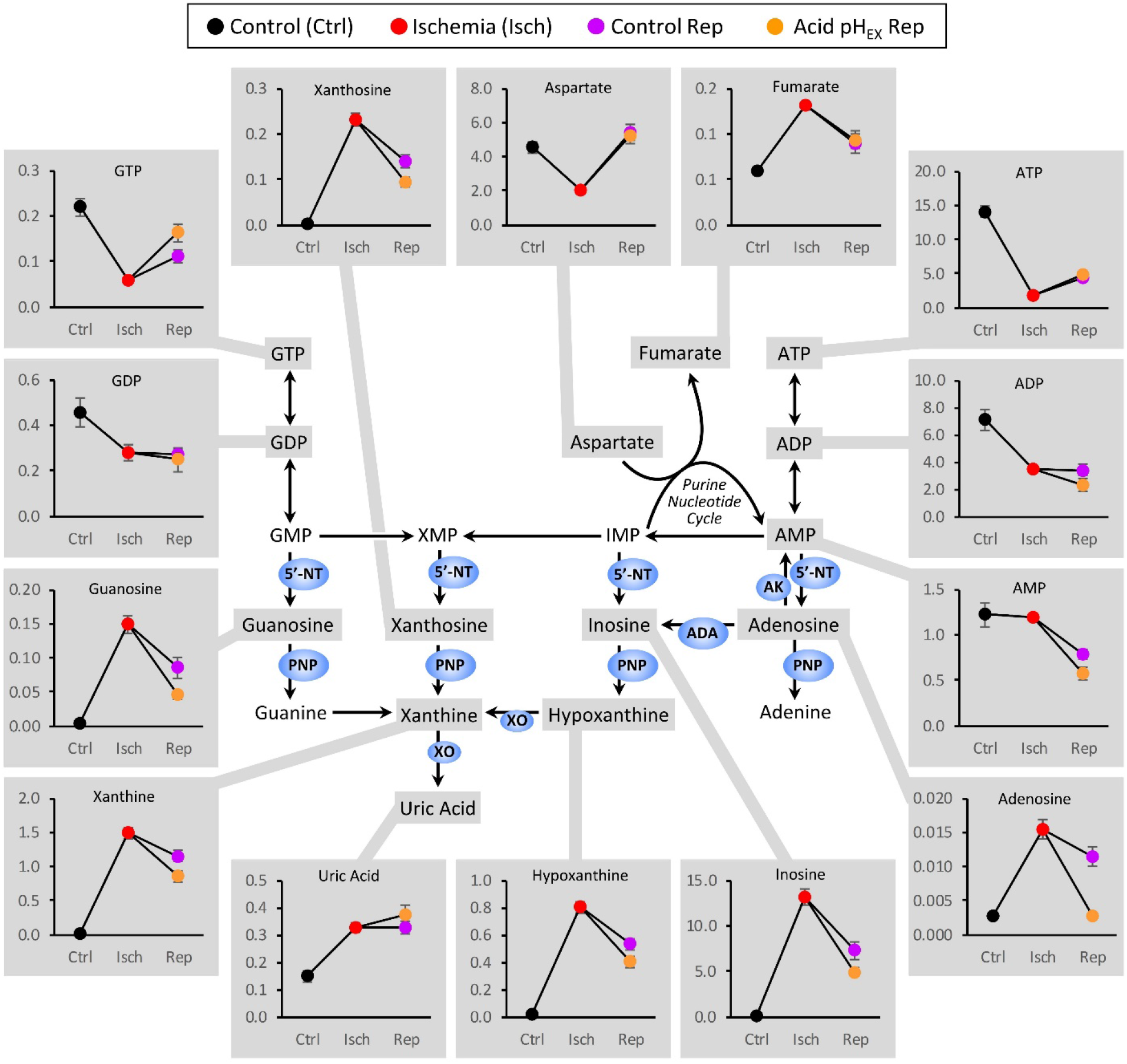
Metabolomics of the purine nucleotide salvage pathway. The pathway is shown with each metabolite that was measured highlighted in gray and linked to its data. Key denotes the color scheme for each treatment group as used in previous figures. Data are means + SEM (N=12-19 per condition) of relative metabolite abundance (arbitrary units).

### 3.7 Impact of acid pH_EX_ reperfusion on metabolite efflux

Upon reperfusion, around ⅓ of accumulated succinate is oxidized to generate ROS, while ⅔ washes out from the heart [15, 24]. Although acidic pH_EX_ has been shown to inhibit succinate release from cells via MCT-1, this occurs at much lower pH_EX_ values (6.0 in [15]) than employed herein (6.8). Furthermore, ischemic succinate that accumulates in the extracellular space will presumably wash out in a manner unaffected by pH.

To examine the impact of acidic pH_EX_ on metabolite release, we measured levels of succinate, fumarate and lactate in effluents collected from hearts during the first 2 minutes of control or acid pH_EX_ reperfusion. As shown in **Figure S6**, acid pH_EX_ reperfusion significantly enhanced succinate efflux during the first minute of reperfusion. This is contrary to the finding that pH_EX_ 6.0 can inhibit succinate release [15], and suggests that mild acidification can enhance succinate efflux. Highlighting the speed with which succinate is oxidized upon reperfusion [23], we also observed that acid pH_EX_ reperfusion enhanced the efflux of fumarate, for which the likely origin is succinate oxidation by mitochondrial complex II. Notably, no impact of pH_EX_ on the efflux of lactate was observed, suggesting the effect on succinate is not a general impact on MCT-1 activity. As such, these data suggest that the cardioprotective benefits of pH_EX_ 6.8 at reperfusion may be mediated in part via enhancing succinate release.

## 4. DISCUSSION

Acidic pH is a hallmark of the enhanced glycolysis that occurs to compensate for impaired mitochondrial Ox-Phos during tissue hypoxia/ischemia. However, surprisingly little is known about the impact of this acidosis on metabolism within the context of hypoxia/ischemia. Furthermore, although extension of ischemic acidosis into early reperfusion is proven to significantly reduce myocardial infarction, the mechanism of this protection is not fully understood, including the role (if any) of altered metabolism.

It is well known that acidosis decreases contractility of cardiac muscle, impacting all stages of excitation-contraction coupling [49–51], including Ca^2+^ dynamics and myofilament function [52–54]. As such, the extension of ischemic acidosis into reperfusion may protect via lowering ATP demand thus allowing cardiomyocytes to re-establish ionic homeostasis.

Maintaining redox homeostasis is also critical for cardiac function and survival, and pH is a known regulator of redox processes [55, 56]. Herein, both the CHIA and A-pH_EX_ studies (**Figures 3 and 5)** revealed that acidic pH caused a shift toward a more reduced intracellular glutathione redox state (higher GSH, lower GSSG). This would be favorable in detoxifying mitochondrial ROS generated at reperfusion [57], and may be a contributing factor that underlies the cardioprotection conferred by acidic pH_EX_ reperfusion.

In considering other potential metabolic differences between control vs. acid pH_EX_ reperfusion that may underlie cardioprotection, the top half of glycolysis was elevated in acid pH_EX_ (**Figure 6**), but this did not appear to be linked to the reactive metabolite MGO, suggesting MGO is not a protective mediator. Likewise, although purine nucleotide salvage appeared to be altered by acid pH_EX_ reperfusion (**Figure 7**), with a potential inhibition of 5NT activity, pharmacologic inhibition of 5NT was unable to recapitulate a protected phenotype. This is possibly because some products of 5NT such as adenosine and inosine are endogenously generated during conditions such as ischemic preconditioning, and are known to confer cardioprotection [58–61]. As such, 5NT inhibition may have a detrimental effect by removing endogenously produced adenosine or inosine from the heart. These findings do not preclude the possibility that some of the protective effect of acid pH_EX_ reperfusion may be mediated by prevention of purine loss.

A surprising finding was the impact of acidic pH_EX_ reperfusion on the release of metabolites from myocardium during the first 2 minutes of reperfusion. Although no impact of pH_EX_ on the release of lactate was seen, there was a significantly larger release of succinate with acid pH_EX_ during the first minute of reperfusion. Previously it has been shown that extreme pH_EX_ acidosis (pH 6.0) can inhibit succinate export via MCT-1, but the current results suggest a complex relationship between pH and succinate release, with mild acidosis causing an increase in release, not a decrease. Such an enhanced release of succinate would be expected to confer a cardioprotective benefit, by lowering the amount of succinate available inside cardiomyocytes for ROS generation [23]. An important caveat is that although more succinate was released from the heart in acid pH_EX_ vs. control reperfusion (**Figure S6**), this did not result in less succinate in the heart itself 2 min. into reperfusion (**Figure 5**). This is likely because succinate is consumed rapidly, such that by the time of sampling, cardiac succinate levels have returned almost to baseline [24]. Sampling at earlier times of reperfusion would be required to track the fate of succinate (both intracellular and extracellular) on a more rapid timescale.

In summary, the CHIA study highlights that acid pH can independently rewire metabolism, although not nearly to the extent that hypoxia or ischemia can. In-fact, there were surprisingly few metabolic alterations unique to acid pH_IN_ that were not also seen with the other two interventions. In contrast the A-pH_EX_ study shows that acid pH_EX_ reperfusion brings about a large number of metabolic changes upon reperfusion, in addition those seen in control reperfusion. At least some of these novel acid-induced metabolic alterations (enhanced succinate release and prevention of purine loss) may partly underlie the cardioprotective benefits of acid pH_EX_ reperfusion. Further studies are required to discern the impact of acid pH_EX_ reperfusion on downstream signaling pathways such as mitochondrial ROS generation and PT pore opening.

## ABBREVIATIONS

5NT: ecto-5’-nucleotidase
A-pH_EX_: Acidic extracellular pH
ATP: adenosine triphosphate
BCECF-AM: 2’-7’-bis-(2-carboxyethyl)-5-(and-6)-carboxyfluorescein, acetoxymethyl ester
CHIA: Control, Hypoxia, Ischemia, Acidic pH_IN_
IR: ischemia-reperfusion
MCT: monocarboxylate transporter
meth-ADP: adenosine 5’-(α,β-methylene) diphosphate
NBC: Na^+^/HCO_3_^-^ cotransporter
NHE: Na^+^/H^+^ exchanger
NMN: Nicotinamide mononucleotide
Ox-Phos: oxidative phosphorylation
pH_EX_: extracellular pH
pH_IN_: intracellular (cytosolic) pH
PT pore: permeability transition pore
ROS: reactive oxygen species
RPP: rate x pressure product
TCA: tricarboxylic acid

## Acknowledgements

We thank Keith Nehrke (Rochester) for use of the imaging system used for BCECF measurements.

## Author Contribution Statement

ASM and PSB designed the research. ASM, JHM, and SMN performed experiments. ASM and PSB analyzed the data and wrote the manuscript. All authors approved the final version of the manuscript.

## SUPPLEMENTAL INFORMATION

### 6 Figures & Legends

**Supplemental Figure 1.**
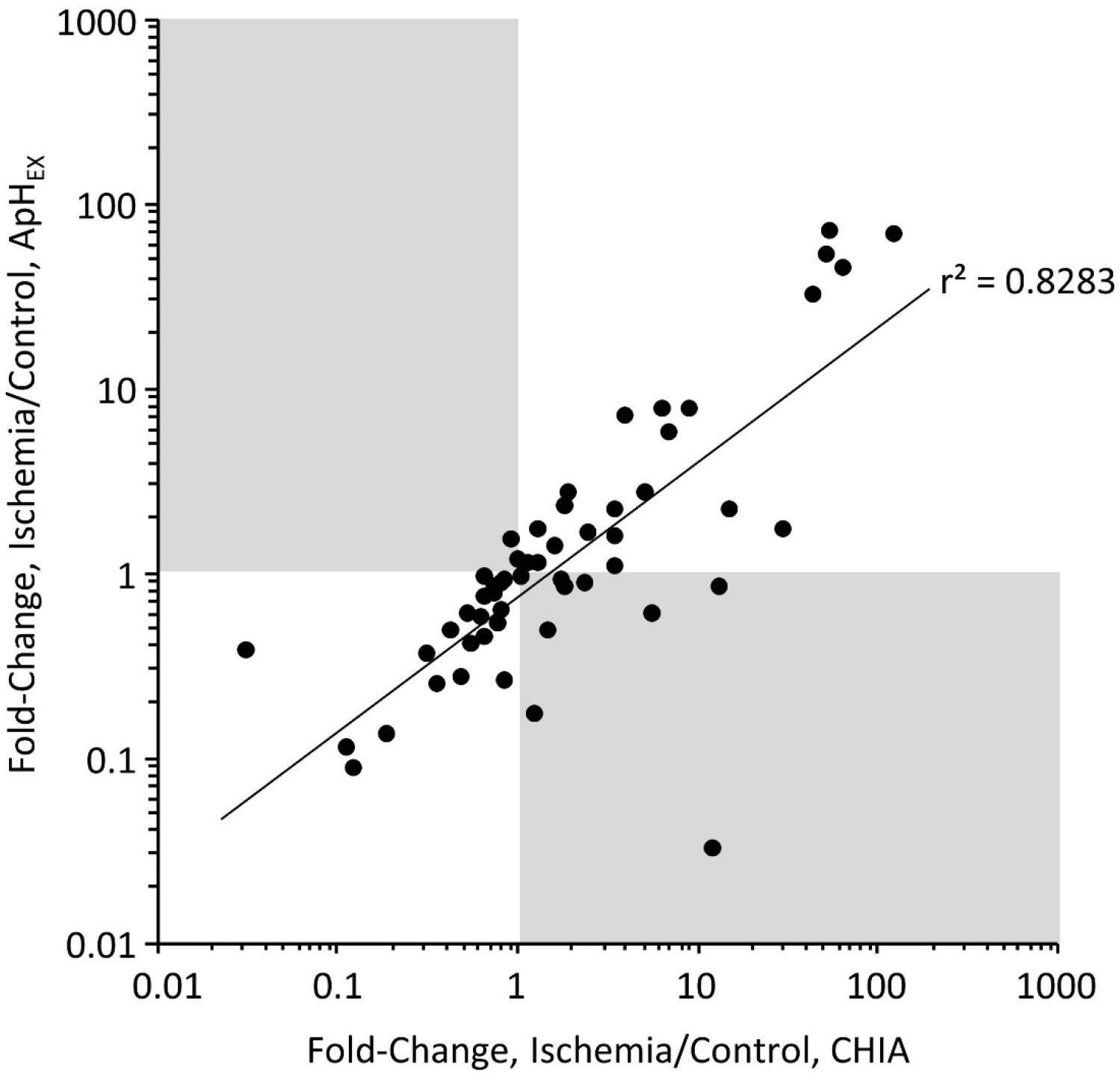
Comparing CHIA and ApH_EX_ ischemic metabolomes. The Log_10_ metabolite fold-changes in ischemia vs. baseline, in the CHIA study (N=8) and ApH_EX_ study (N=19). Correlation coefficient, r^2^ = 0.8283.

**Supplemental Figure 2.**
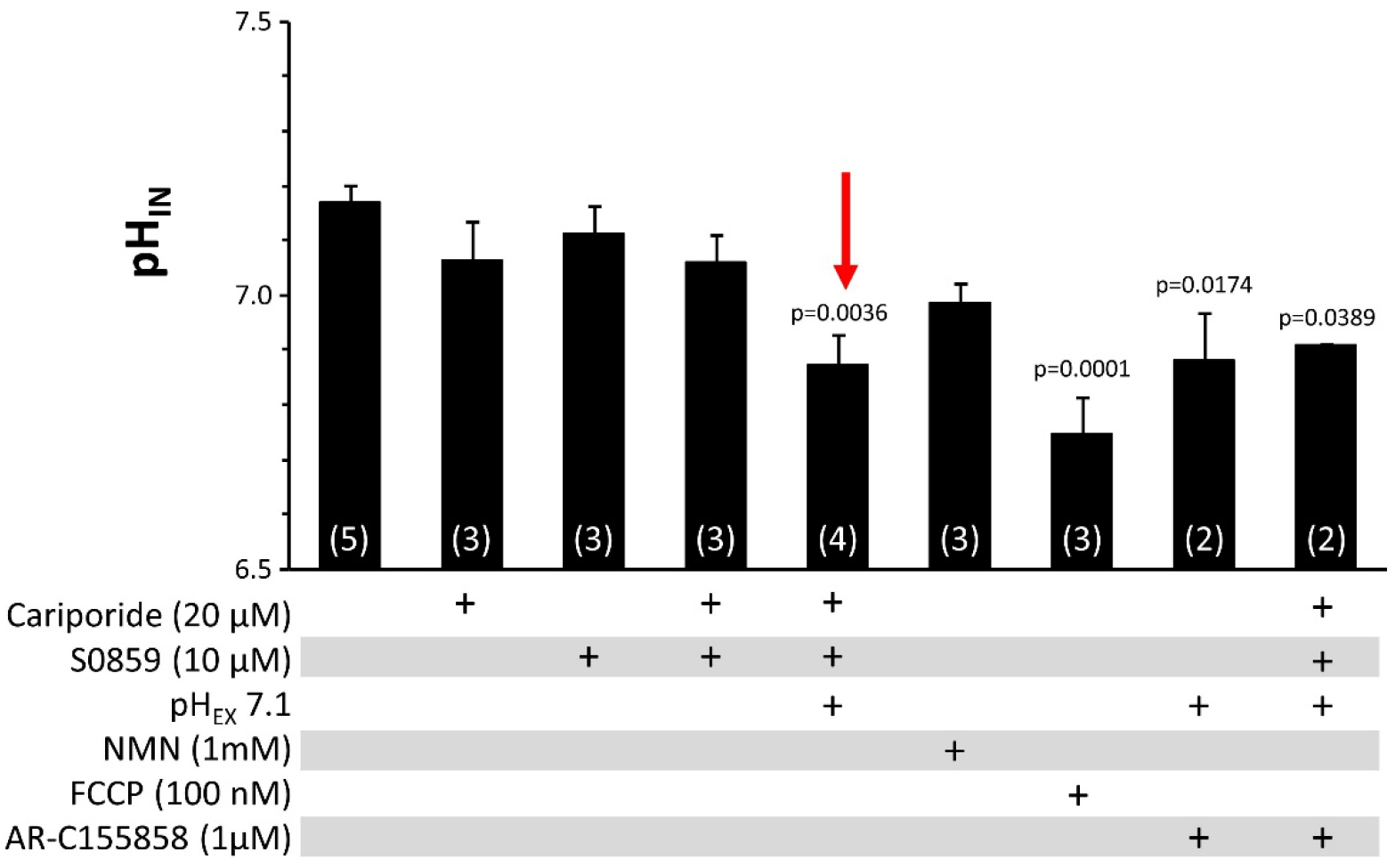
Screening conditions for cardiomyocyte pH_IN_ acidification. Various physiologic and pharmacologic interventions (concentrations as noted) were screened to acidify isolated mouse cardiomyocytes. Intracellular pH (pH_IN_) was measured using BCECF-AM. Data are means ± SEM (N for each condition shown on graph bars). p-values (ANOVA followed by Tukey HSD test) that are < 0.05 are denoted above error bars. Red arrow indicates the condition used for acid pH_IN_ for the CHIA study.

**Supplemental Figure 3.**
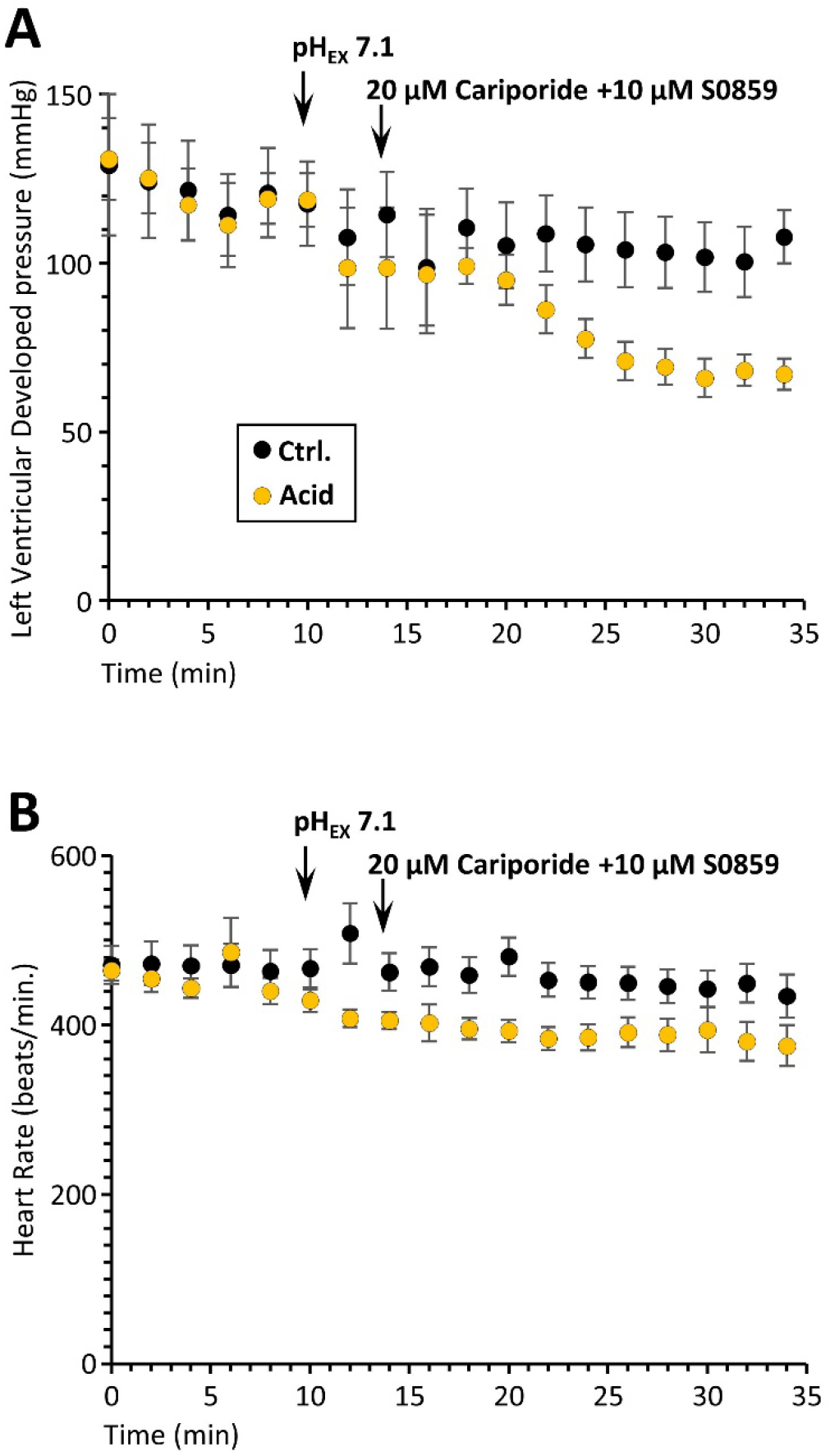
Impact of acid pH_IN_ on rate pressure product (RPP) components. Graphs show **(A):** left ventricular developed pressure (LVDP, mmHg) and **(B):** heart rate (beats/min), following imposition of acid pH_IN_ conditions as indicated by arrows: Buffer change (pH_EX_ 7.1) and drug administration (cariporide + S09859). Data are means ± SEM (N=5-7).

**Supplemental Figure 4.**
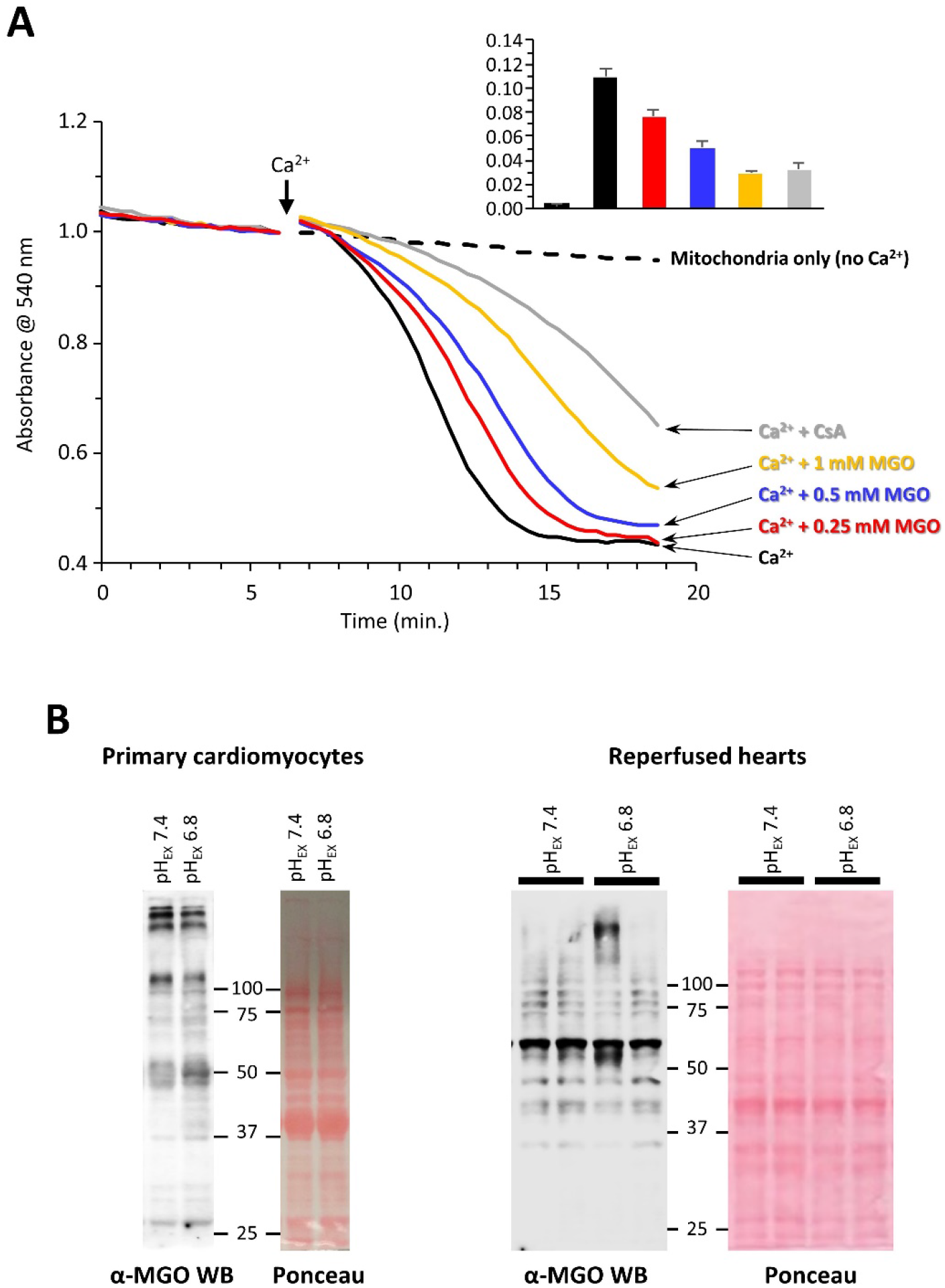
MGO inhibits mitochondrial PT pore opening. **(A):** Opening of the mitochondrial PT pore was assayed spectrophotometrically at 540nm in isolated mitochondria. Example traces (representative of N=3) are shown, with addition of 100 μM CaCl_2_ to initiate PT pore opening indicated by the arrow. Traces are color coded denoting concentrations of methylglyoxal (MGO, 0.25 – 1 mM) added, or the PT pore inhibitor cyclosporin A (CsA, gray). Dashed line is no Ca^2+^ control. **Inset:** Rate of swelling, means from N=3 ± SEM. **(B/C)**: Western blots showing abundance of MGO protein adducts in **(B):** primary cardiomyocytes (N=1), or **(C):** reperfused mouse hearts, subject to either control pH_EX_ 7.4 or acid pH_EX_ 6.8. Ponceau stained membranes to indicate equal protein loading are shown adjacent.

**Supplemental Figure 5.**
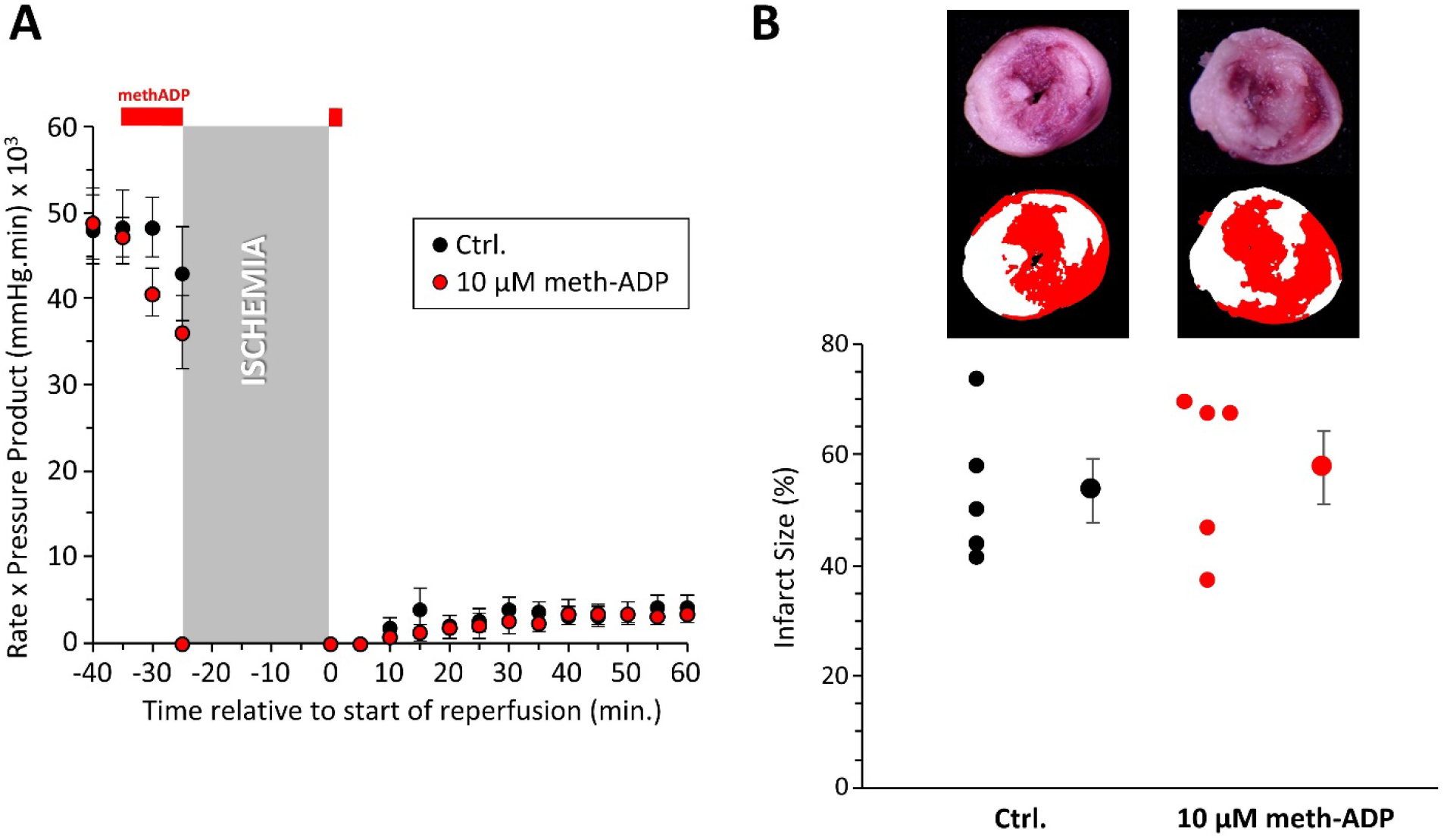
Impact of inhibiting ecto-5’-NT with methylene ADP on IR injury. **(A):** Cardiac function (rate x pressure product, RPP) during IR. The 5’-NT inhibitor, methylene ADP (10 μM, meth-ADP) was infused pre-ischemically for 10 min. and for 2 min. into reperfusion, as shown in red above the graph. Data are means ± SEM (N=6). **(B):** Post IR triphenyltetrazolium chloride-stained heart cross-sections for control and meth-ADP. Representative heart slices are shown above the graph with pseudo-colored mask (white = infarct, red = live tissue) used for planimetry. Graph shows individual data points on the left, means ± SEM on the right (N for each condition seen from number of data points).

**Supplemental Figure 6.**
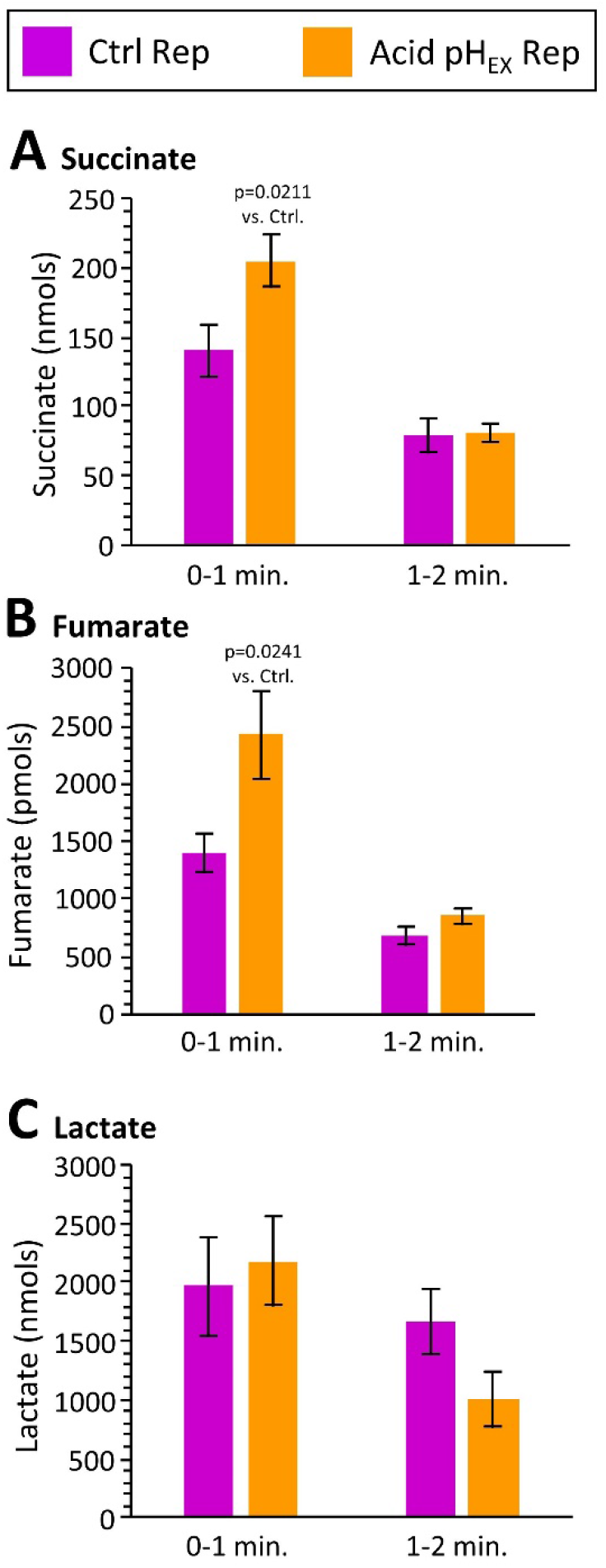
HPLC quantitation of metabolites in cardiac effluent. Cardiac perfusates (effluents) were collected in 1 min. intervals for the first 2 min. of reperfusion, from control or acid pH_EX_ reperfused hearts. Perfusion at 4 ml/min. yielded 4 ml of perfusate. Metabolites were quantified by HPLC. Data for **(A):** succinate, **(B):** fumarate, **(C):** lactate are shown as nmols per 4ml of perfusate, and are means ± SEM, N=14-16. p-values above bars (Tukey’s HSD test) denote significance between control vs. acid pH_EX_ REP groups (non-significant values are not shown). Lactate data are corrected for presence of 1.2 mM lactate in KHB.

